# Inhibition of Biofilm Formation by Modified Oxylipins from the Shipworm Symbiont *Teredinibacter turnerae*

**DOI:** 10.1101/2020.11.22.393314

**Authors:** Noel M. Lacerna, Cydee Marie V. Ramones, Jose Miguel D. Robes, Myra Ruth D. Picart, Jortan O. Tun, Bailey W. Miller, Margo G. Haygood, Eric W. Schmidt, Lilibeth A. Salvador-Reyes, Gisela P. Concepcion

**Affiliations:** The Marine Science Institute, University of the Philippines Diliman, Quezon City 1101, Philippines; Department of Medicinal Chemistry, University of Utah, Salt Lake City, Utah 84112, United States

**Keywords:** antibiofilm, oxylipins, *Staphylococcus epidermidis*, *Lyrodus*, shipworm, *Teredinibacter turnerae*

## Abstract

Bioactivity-guided purification of the culture broth of the shipworm endosymbiont *Teredinibacter turnerae* 991H.S.0a.06 yielded a new fatty acid, turneroic acid (**1**), and two previously described oxylipins (**2-3**). Turneroic acid (**1**) is an 18-carbon fatty acid decorated by a hydroxy group and an epoxide ring. Compounds **1-3** inhibited bacterial biofilm formation in *Staphylococcus epidermidis*, while only **3** showed antimicrobial activity against planktonic *S. epidermidis*. Comparison of the bioactivity of **1-3** with structurally related compounds indicated the importance of the epoxide moiety for selective and potent biofilm inhibition.

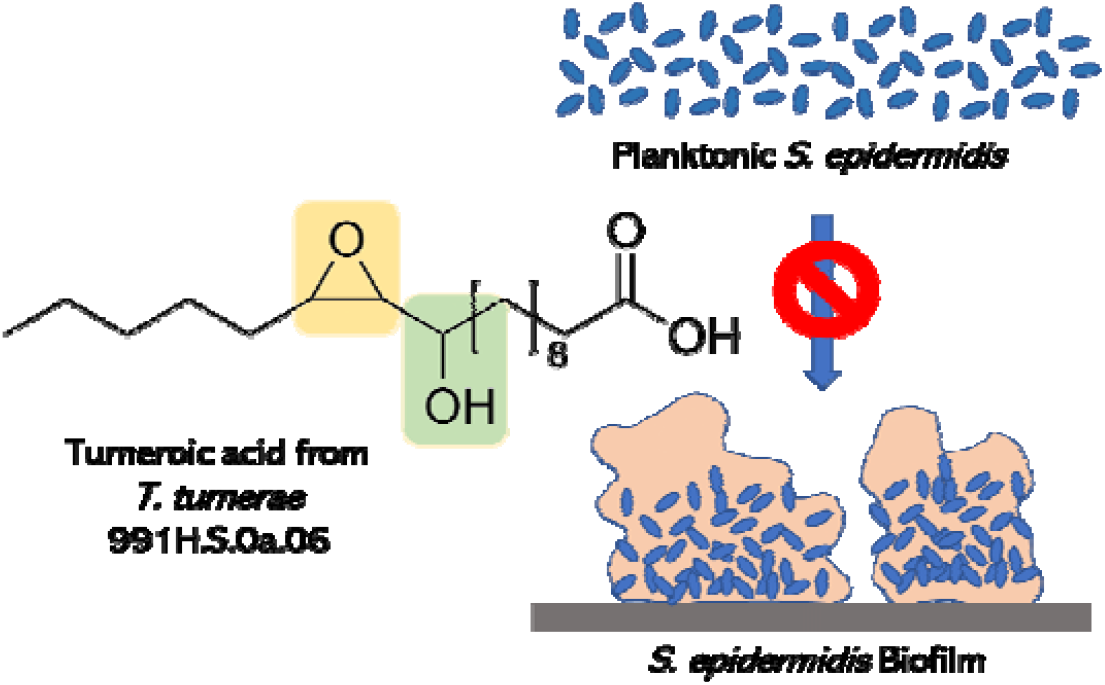

## 1. Introduction

Microorganisms often colonize surfaces by creating a protected and nutritionally rich ecological niche called a biofilm [1]. Biofilms provide protected access to nutritional sources, facilitate cellular communication, and increase resistance to antimicrobials in concentrations normally lethal to planktonic organisms [2,3]. *S. epidermidis*, a normally commensal bacterium common in skin and mucous membranes that is pathogenic in certain circumstances, is considered the best-known example of a pathogen that uses biofilm formation as a virulence factor [4]. *S. epidermidis* biofilm-mediated infections represent the most common infections on indwelling medical devices and the only effective treatment is replacement of an infected device with new and uninfected ones [4,5]. Currently, research is focused on understanding the in-depth mechanism of biofilm formation. The use of small molecules affecting biofilm-associated infections could possibly provide a solution to this longstanding problem.

Nature has historically been the most productive source of antibiotics [6]. In recent years, a strategy has been to search for bioactive compounds in previously overlooked habitats such as host-associated bacteria [7,8]. Shipworms (family: *Teredinidae*) are marine wood-boring bivalves that possess bacterial endosymbionts in bacteriocytes located in the gills [9,10]. The specialized intracellular bacteria support the host’s nutrition by producing enzymes used to digest cellulose and fix nitrogen [11]. One of the major bacterial symbionts in shipworms is *Teredinibacter turnerae*, although several other species have also been identified. Genomic analysis of *T. turnerae* strain T7901 showed that an unusually high proportion of the genome (7%) is devoted to secondary metabolism [12], comparable to that of the biomedically-important Actinobacteria. The substantial investment of *T. turnerae* in secondary metabolite biosynthesis suggests a potential role for small molecules in the symbiotic relationship between *T. turnerae* and shipworm host as well as other microorganisms. For example, the macrodiolide antibiotics tartrolons produced by *T. turnerae* T7901 were shown to inhibit other shipworm symbionts and free-living pathogenic bacteria [13]. The siderophore turnerbactins provide another mechanism for microbial competition in the host through iron sequestration [9]. However, despite the vast potential for natural product discovery highlighted by the genomic analysis, few compounds have been isolated and characterized from this interesting bacterial group.

In this study, we screened shipworm-associated microorganisms for antimicrobial and biofilm inhibitory activity. One of the bioactive isolates, *Teredinibacter turnerae* 991H.S.0a.06 from *Lyrodus pedicellatus* was subjected to bioassay-guided fractionation, affording the new compound, turneroic acid (**1**), and two known fatty acids (**2-3**). Structure-activity comparison of **1-3** with related compounds provided insights on the critical pharmacophore for potent and selective biofilm inhibition.

## 2. Results and Discussion

### 2.1 Structure Elucidation

Turneroic acid (**1**) was purified as a white powder with λ_max_ of 230 nm. HRESIMS of **1** showed a sodiated peak at *m/z* 337.2363, which was assigned to a molecular formula of C_18_H_34_O_4_Na. The ^1^H NMR profile of turneroic acid (**1**) showed characteristic signals for oxylipins with a terminal methyl (δ_H_ 0.92), methylene envelope (δ_H_1.33-1.35) and α-methylenes (δ_H_ 2.24). COSY, HSQC and HMBC showed distinctive signals for a carboxylic acid (δ_C_ 178.1), an epoxide moiety (δ_C_/δ_H_ 63.4/2.67; 57.7/2.86) and a hydroxylated methine (δ_C_/δ_H_ 72.8/3.30). 2D NMR correlations allowed for the assignment of three partial structures (Figure 1a). What remained after the assignment of these partial structures was a methylene envelope consisting of a C5H10 unit, in which the ^1^H and ^13^C shifts were overlapping. While minor impurities were observed in the TOCSY spectrum of **1** due to prolonged storage in solution, clear correlations were observed between H-13/H-18 and H-14/H-18 and established the linkage between partial structures I and II. This was corroborated by the lack of TOCSY correlation between the terminal CH_3_ (δ_H_ 0.92) and the hydroxylated methine (δ_H_ 3.30). HMBC correlations between H-2/C-4, H-9/C-8, H-9/C-7 and H-10/C-8 indicated that partial structures II and III are linked by the methylene envelope. Hence, the structure of **1** was assigned as 11-hydroxy-12,13-epoxy-octadecanoic acid. A somewhat similar compound, except with additional unsaturation, was previously synthesized, although no spectroscopic or configurational data were provided for the synthetic compound. Since the synthetic compound potentially consists of a mixture of 4 diastereomers as drawn, the identity of the compound is somewhat unclear [14].

**Figure 1.**
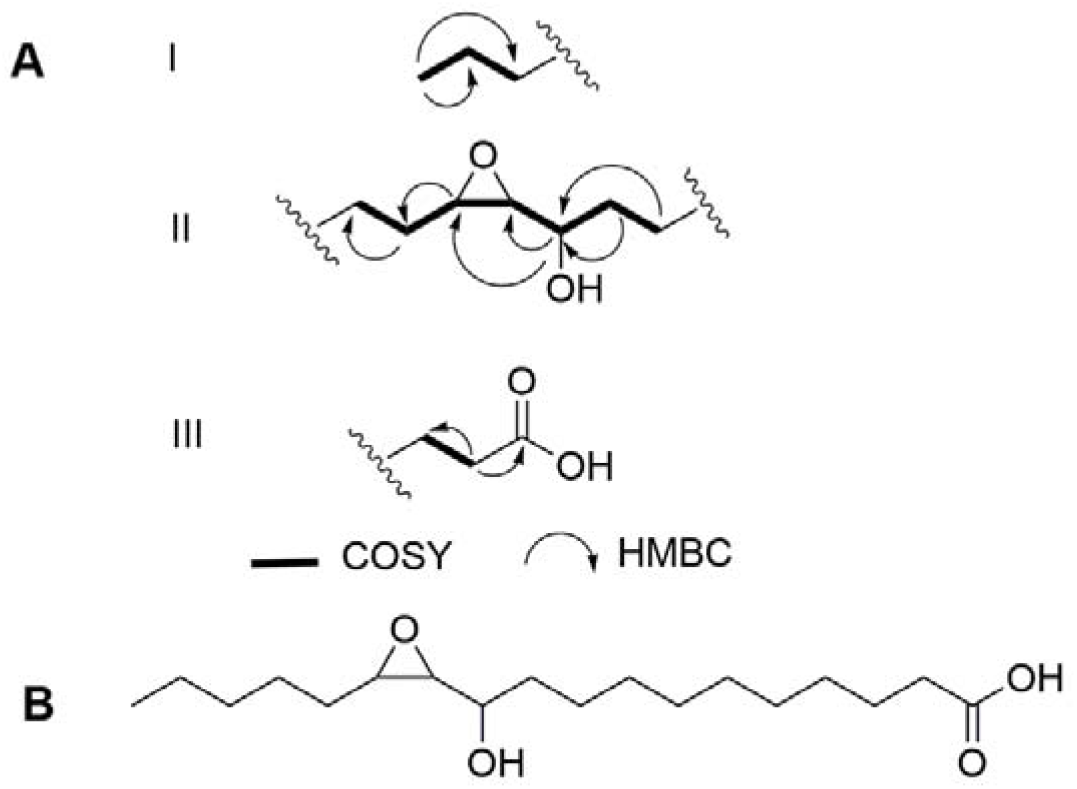
a.) Key fragments with COSY and HMBC correlations and b.) structure of turneroic acid (**1**).

The ^1^H NMR spectrum of **2** and **3** also showed characteristic signals for fatty acids (Fig. S11 and S12, Supporting Information). The ^1^H NMR and MS data for **2** matched the known compound (*E*)-9- oxohexadec-10-enoic acid previously purified from the red alga *Gracilaria verrucosa* (Figure 2) [15]. The ^1^H NMR data for **3** showed marked similarity with **2**, indicating that these are analogues. HRESIMS analysis of **3** gave a 28 Da mass difference with **2**, suggesting two additional –CH_2_s in **3**. This is in accordance with the additional methylene protons observed in the ^1^H NMR spectrum of **3** (Fig. S12, Supporting Information). Dereplication and further MS/MS fragmentation indicated that **3** is (*E*)-11-oxooctadec-12-enoic acid (Figure 2), previously reported from the marine green alga *Ulva fasciata* [16] (Fig. S10, Supporting Information).

**Figure 2.**
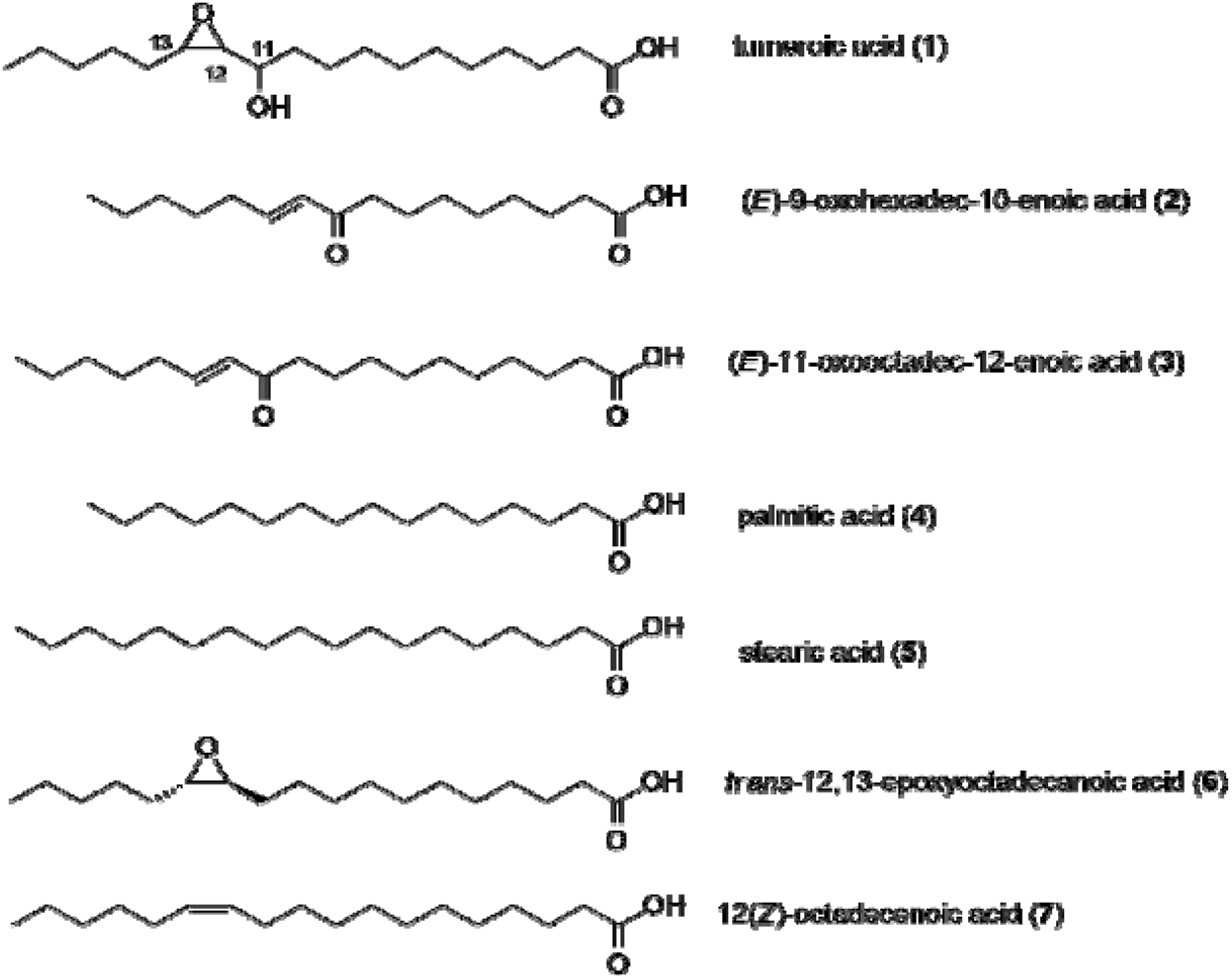
Structure of modified oxylipins.

Compounds **1** and **3** are structurally related. Both are 18-carbon fatty acids, with structural modifications at C11-C13. Turneroic acid bears an epoxide at C12-C13 instead of a C=C as in **3** and a hydroxy instead of a keto group at C-11. This may likely point to turneroic acid being a biotransformation product of **3**. This is analogous to the enzymatic and non-enzymatic conversion of linoleic acid to the corresponding hydroxyepoxy-octadecenoic acid in human plasma [14]. On the basis of the same biosynthetic origin for **1** and **3**, the relative configuration of the epoxide moiety in **1** is likely to be *trans*. The ^1^H NMR spectrum of **3** gave a coupling constant of *J ∼*15.7 Hz for the olefinic protons (δ_H_ = 6.12 and 6.95), suggesting a *trans* coupling between H-12 and H-13. The configuration of turneroic acid (**1**) could not be assigned due to insufficient material. Nonetheless, the structural features inspired us to explore biological activity using simpler analogues (see below).

### 2.2 Biological Activities of Oxylipins (**1-7**)

Compounds **1-3** are structurally related to the naturally occurring fatty acids palmitic acid (**4**) and stearic acid (**5**), as well as possible biotransformation compounds *trans*-12,13-epoxy-octadecanoic acid (**6**) and 12(*Z*)-octadecenoic acid (**7**) (Figure 2). All seven compounds were subjected to a panel of bioactivity assays to assess their effects on biofilm formation, planktonic microbial pathogen proliferation, and mammalian cell line cytotoxicity (Table 2). Not surprisingly, the saturated C16 and C18 fatty acids, palmitic (**4**) and stearic acids (**5**), respectively, did not show any significant bioactivity against microbial pathogens or mammalian cells (Table 2). Only **3** was bioactive against planktonic *S. epidermidis* (MIC = 24.5 *µ*g/mL), while **7** showed weak inhibition against planktonic *S. aureus* (MIC = 64.0 *µ*g/mL). Four of the seven compounds (**1, 2, 3**, and **6**) inhibited biofilm formation in *S. epidermidis* (Table 2). Compound **3** demonstrated cytotoxicity in mammalian MDCK cells.

**Table 2.** Minimum inhibitory concentration (MIC) of **1-7** against biofilm formation of *S. epidermidis* RP62A ATCC 35984^™^ and planktonic cells of *S. epidermidis* RP62A ATCC 35984™, *S. aureus* ATCC 6538^™^ and methicillin-resistant *S. aureus* ATCC 43300^™^.^*a*^

| Compounds | MIC (µg/mL)<br>*S. epidermidis*<br>RP62A (ATCC35984) |  | Selectivity<br>Index<br>(MIC<sub>Planktonic</sub>/<br>MIC<sub>Biofilm</sub>) | MIC (µg/mL)<br>*S. aureus*<br>(ATCC6538) | MIC (µg/mL)<br>methicillin-resistant<br>*S. aureus*<br>(ATCC43300) | IC<sub>50</sub> (µg/mL)<br>MDCK NBL-2<br>(ATCC CCL-34) |
| --- | --- | --- | --- | --- | --- | --- |
|  | Biofilm | Planktonic |  |  |  |  |
| **1** | 32.0 | 128 | 4 | >128 | >128 | 61.7 |
| **2** | 90.0 | >128 | >1.4 | >128 | >128 | 33.0 |
| **3** | 11.2 | 24.5 | 2.2 | >128 | >128 | 13.6 |
| **4** | >128 | >128 | ~1 | >128 | >128 | >128 |
| **5** | >128 | >128 | ~1 | >128 | >128 | >128 |
| **6** | 16.0 | >128 | >8 | >128 | >128 | >128 |
| **7** | >128 | > 128 | ~1 | 64.0 | >128 | >128 |
| dispersin B | 50.0 | >>500 | >>10 | n/a | n/a | n/a |
| chloramphenicol | n/a | 8 | n/a | n/a | n/a | n/a |
| oxacillin | n/a | n/a | n/a | 8 | 1 | n/a |

<sup>a</sup> Data presented as MIC of two independent trials performed in quadruplicates. n/a = not applicable.

A comparison of each structure to its bioactivity profile shows that decoration of the hydrophobic chain with various functional groups leads to a variety of biological activities. For example, in the C16 fatty acids **2** and **4**, the addition of an α, β-unsaturation in **4** leads to cytotoxic activity against mammalian cells and modest inhibition of biofilm formation in *S. epidermidis* (Table 2). Comparing **2** and **3**, the addition of one CH_2_ group in the latter resulted in potent but non-selective activity against biofilm, planktonic *S. epidermidis*, and MDCK cells. Improved selectivity for biofilm inhibition was observed for compounds bearing an epoxide ring, as in **1** and **6**. The selectivity index increased by 2- to 4-fold with the introduction of the epoxide functionality. At the same time, **6** showed no antiproliferative action in mammalian cells.

The potency and selectivity of the oxylipins are dictated by the addition of functional groups to the lipophilic chain. Having an electrophilic center improves the potency, while the nature of the functional groups tunes the selectivity. Electrophilic centers are a common feature among other compounds with biofilm inhibitory activity such as in *cis*-2-decenoic acid [17,18], cinnamaldehyde [19], pentadecanal [20], 5-dodecanolide [21], parthenolide [22] and cembranoid alkaloid [23]. Pentadecanal, from the marine bacterium *Pseudoalteromonas haloplanktis* TAC125, is structurally related to the *Vibrio harveyi* quorum sensing molecule tetradecanal.[20]. 5-dodecanolide restricts biofilm formation in MRSA by decreasing the expression of the quorum sensing genes *agrA* and *agrC*, which consequently downregulates the expression of effector genes involved in biofilm formation [21]. The transcriptional effects of 5-dodecanolide is suggested to affect the early stages of biofilm formation by modulating the levels of adhesion proteins [21]. The anti-biofilm compounds **1-3** and **6** are similar to reactive electrophilic oxylipins which possess qualities of being lipophilic and thiol reactive. Thiol-modification is assumed to represent a key mechanism in signal transduction [24]. These oxylipins chemically modify several proteins particularly thioredoxins which function in redox signaling and oxidative stress responses. A thioredoxin *trxH*_*Ep*,_ from a fish pathogen *Edwardsiella piscicida*, was identified to contribute to the microorganism’s adversity adaptation and its pathogenicity (25). Deletion of *trxH*_*Ep*,_ led to retarded bacterial biofilm growth [25]. Because the anti-biofilm activity is correlated with the presence of thiol-reactive groups, it is possible that **1-3** and **6** similarly function by covalently modifying bacterial proteins. Further work is required to test this hypothesis.

The existence of anti-biofilm compounds such as **1-3** from *T. turnerae* may have an ecological relevance to the shipworm host. Compounds **1-3** could possibly be part of an arsenal of compounds, including the previously reported tartrolons and turnerbactins, for suppressing microbial growth within the caecum of the shipworm host, where lignocellulose is degraded, and glucose is liberated. This would constitute a complex strategy involving inhibition of biofilms with **1**- **3**, sequestration of iron by turnerbactin, and planktonic cell killing by the tartrolons in order to limit the proliferation of competing microbes [9,13]. Furthermore, the biotransformation of fatty acids to oxylipins by *T. turnerae* highlights an effective strategy to modulate the potency and selectivity of biological activity, starting with simple and abundant fatty acid precursors.

## 3. Materials and Methods

### 3.1 General Experimental Procedures

^1^H, ^13^C and 2D NMR data were collected using a Varian Inova 500 MHz spectrometer equipped with a 3 mm NMR probe using residual solvent signals for referencing. Low resolution ESI-MS was done using Shimadzu 8040 through direct injection in 50% MeOH:H_2_O with 0.1% formic acid. High resolution MS (HRMS) data was obtained using Waters Xevo G2 XS QTOF through direct infusion using 50% aqueous ACN with 0.1% formic acid. HPLC purification was done using Shimadzu High Performance Liquid Chromatography (HPLC) equipped with binary pumps, fraction collector and photodiode array detector (Shimadzu Kyoto, Japan). Fatty acids palmitic (**4**) and stearic acids (**5**) were sourced from Sigma-Aldrich (St. Louis, MO, USA). (±) *trans*-12,13-Epoxy-octadecanoic acid (**6**) and 12(*Z*)-octadecenoic acid (**7**) were purchased from Larodan Fine Chemicals (Solna, Sweden).

### 3.2 Biological Material

*T. turnerae* strain 991H.S.0a.06 was obtained from the gill of *Lyrodus pedicellatus* collected in Panglao, Bohol and was identified through amplification and sequencing of 16s rRNA genes using primers 27F (5’-AGAGTTTGATCMTGGCTCAG-3’) and 1492R (5’- TACGGYTACCTTGTTACGACTT-3’). Sequences were submitted to GenBank with accession number MH379668.

### 3.3 Culturing and Extraction

*T. turnerae* 991H.S.0a.06 was grown for 7 days using Shipworm Basal Medium broth at 30 °C, 250 rpm. SBM contained sucrose (5 g/L), NaCl (19.8 g/L), NH_4_Cl (267.5 mg/L), MgCl_2_·6H_2_O (8.95 g/L), Na_2_SO_4_ (3.31 g/L), CaCl_2_·2H_2_O (1.25 g/L), NaHCO_3_ (0.162 g/L), Na_2_CO_3_ (10 mg/L), KCl (0.552 gm/L), KBr (81 mg/L), H_3_BO_3_ (21.5 mg/L), SrCl_2_·6H_2_O (19.8 mg/L), KH_2_PO_4_ (3.82 mg/L), NaF (2.48 mg/L), Na_2_MoO_4_·2H_2_O (2.5 mg/L), MnCl_2_·4H_2_O (1.8 mg/ L), ZnSO_4_.7H_2_O (0.22 mg/L), CuSO_4_·5H_2_O (0.079 mg/L), Co(NO_3_)_2_·6H_2_O (0.049 mg/L), Fe-EDTA complex (4.15 mg/L), and HEPES (4.76 g/L) adjusted to pH = 8.0). After the incubation period, the broth was centrifuged and the supernatant was extracted with HP20^™^ Diaion resin for 2 h. The filtered resin was washed with H_2_O, 75% aqueous MeOH then eluted with 100% MeOH. The methanolic extract was dried under reduced pressure to yield the crude extract. This was subjected to solvent partitioning (3x) with EtOAc:H_2_O (1:1 v/v). The organic extract was separated into four fractions using reversed phase C18 column chromatography with a step-gradient elution of MeOH in H_2_O (40%, 60%, 80% and 100%). The bioactive 100% MeOH C18 open column fraction was further separated by size exclusion chromatography (Sephadex LH20) using MeOH as eluting solvent. A total of 16 LH20 fractions were collected and subjected to silica thin layer chromatography (TLC) using a 9:1 CH_2_Cl_2_:MeOH system as developing solvent. Fractions were pooled based on similarity of the TLC profile to generate four fractions (Fractions 1-4). LH20 Fraction 2 was subjected to semipreparative reversed phase HPLC (Varian Polaris C-18 A-50, 250 x 10 mm, 5 µm; flowrate: 3.00 mL/min) in a linear gradient of MeOH:H_2_O with 0.1% TFA as solvent system (60% MeOH:H_2_O for 5 min, 60%-100% MeOH:H_2_O over 7.5 min, 100% MeOH for 5 min). Fractions were collected at 1 min interval using a fraction collector. The bioactive fraction with t_R_= 15.0 min, exhibiting antibiofilm activity, was further purified by semipreparative HPLC (Phenomenex Luna 5µ C18, 5µm; 250 x 10 mm; flow rate, 3 mL/min) using a linear gradient of MeOH:H_2_O with 0.1% TFA (75% MeOH:H_2_O for 5 min, 75-95% MeOH:H_2_O over 5 min, 100% MeOH for 15 min) to afford turneroic acid (**1**, 1.4 mg, t_R_ = 14.7 min). LH20 Fraction 4 was also fractionated using semipreparative RP-HPLC (Varian Polaris C-18 A-50, 250 x 10 mm, 5 µm; flowrate: 3 mL/min) with MeOH:H_2_O with 0.1% TFA as solvent system (75- 100% MeOH:H_2_O in 25 min, 100% MeOH for 5 min) to afford **2** (3.2 mg, t_R_= 13.5 min) and **3** (6.5 mg, t_R_ = 17.4 min).

#### Turneroic acid/11-hydroxy-12,13-epoxyoctadecanoic acid (**1**)

white powder; UV (MeOH + 0.1% TFA) λ_max_230 nm; LC-ESIMS m/z 315.40 [M+H]^+^;HRESIMS *m/z* 337. 2363 [M + Na]^+^ (calcd for C_18_H_34_O_4_Na 337.2355); ^1^H NMR and ^13^C NMR data, see Table 1.

**Table 1.**
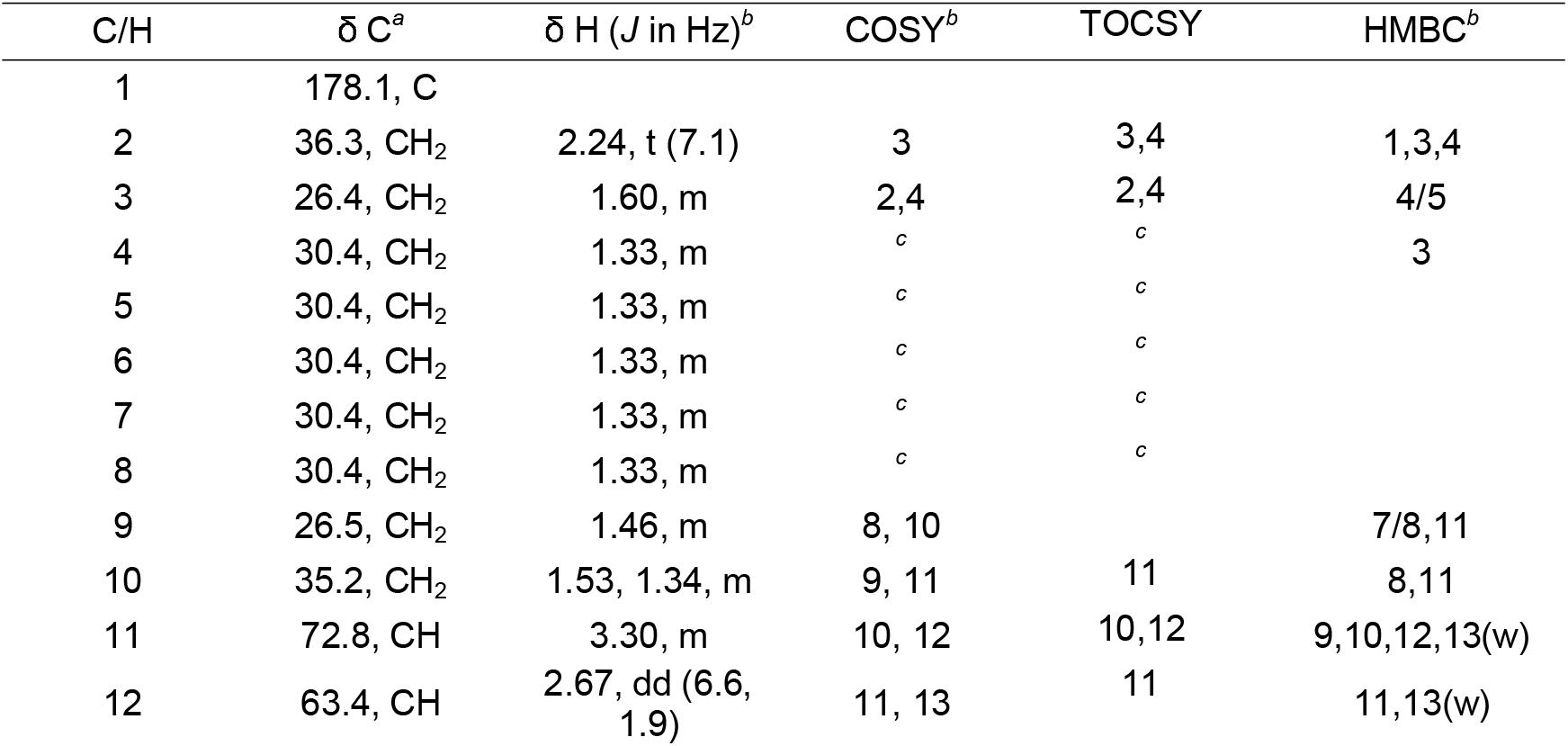
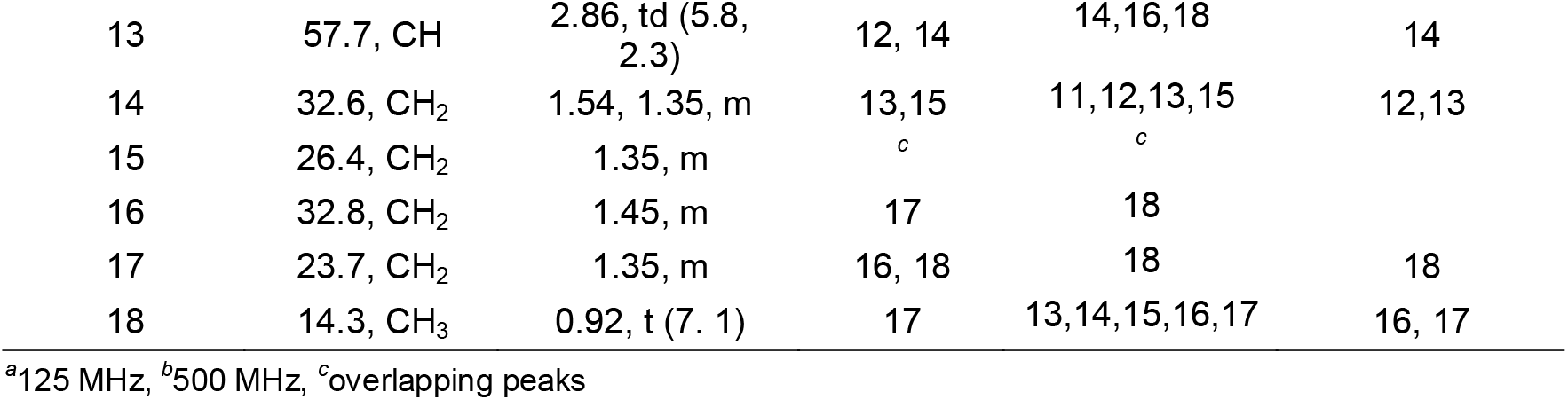
NMR Spectroscopic Data for Turneroic Acid (**1**) in CD_3_OD.

| C/H | $\delta \text{ C}^a$ | $\delta \text{ H (J in Hz)}^b$ | COSY<sup>b</sup> | TOCSY | HMBC<sup>b</sup> |
| --- | --- | --- | --- | --- | --- |
| 1 | 178.1, C |  |  |  |  |
| 2 | 36.3, $\text{CH}_2$ | 2.24, t (7.1) | 3 | 3,4 | 1,3,4 |
| 3 | 26.4, $\text{CH}_2$ | 1.60, m | 2,4 | 2,4 | 4/5 |
| 4 | 30.4, $\text{CH}_2$ | 1.33, m | <sup>c</sup> | <sup>c</sup> | 3 |
| 5 | 30.4, $\text{CH}_2$ | 1.33, m | <sup>c</sup> | <sup>c</sup> | |
| 6 | 30.4, $\text{CH}_2$ | 1.33, m | <sup>c</sup> | <sup>c</sup> | |
| 7 | 30.4, $\text{CH}_2$ | 1.33, m | <sup>c</sup> | <sup>c</sup> | |
| 8 | 30.4, $\text{CH}_2$ | 1.33, m | <sup>c</sup> | <sup>c</sup> | |
| 9 | 26.5, $\text{CH}_2$ | 1.46, m | 8, 10 | | 7/8,11 |
| 10 | 35.2, $\text{CH}_2$ | 1.53, 1.34, m | 9, 11 | 11 | 8,11 |
| 11 | 72.8, CH | 3.30, m | 10, 12 | 10,12 | 9,10,12,13(w) |
| 12 | 63.4, CH | 2.67, dd (6.6, 1.9) | 11, 13 | 11 | 11,13(w) |
| 13 | 57.7, CH | 2.86, td (5.8, 2.3) | 12, 14 | 14,16,18 | 14 |
| 14 | 32.6, CH<sub>2</sub> | 1.54, 1.35, m | 13,15 | 11,12,13,15 | 12,13 |
| 15 | 26.4, CH<sub>2</sub> | 1.35, m | <sup>c</sup> | <sup>c</sup> |  |
| 16 | 32.8, CH<sub>2</sub> | 1.45, m | 17 | 18 |  |
| 17 | 23.7, CH<sub>2</sub> | 1.35, m | 16, 18 | 18 | 18 |
| 18 | 14.3, CH<sub>3</sub> | 0.92, t (7. 1) | 17 | 13,14,15,16,17 | 16, 17 |

<sup>a</sup>125 MHz, <sup>b</sup>500 MHz, <sup>c</sup>overlapping peaks

### 3.4 Antimicrobial Microdilution Assay

A glycerol stock of *Staphylococcus aureus (*ATCC6538) was thawed and streaked on Mueller Hinton Agar (MHA) plates. Methicillin-resistant *S. aureus* (MRSA) (ATCC 43300) was streaked on MHA supplemented with oxacillin (8 *µ*g/mL). Plates were incubated at 37 °C for 8-12 h. Colonies from the MHA plate were inoculated Mueller Hinton broth (MHB) (7 mL) with 2% NaCl and incubated for 6 h, 37 °C, with shaking at 150 rpm. The broth culture was adjusted according to 0.5 McFarland standard (1 x 10^8^cells/mL) and was further diluted 200-fold and used as the final inoculum for the assay. The test organism (200 *µ*L) was dispensed to each well of a 96-well plate. Compounds **1-7**, positive control and 1% DMSO as solvent control were tested using a two-fold dilution scheme starting at 0.158 *µ*g/mL with ten dilutions each. Finally, 0.02% resazurin (20 *µ*L) was added into each well after 24 h and the fluorescence signal was measured using 530 nm excitation and 590 nm emission filters using a Biotek Synergy HT microplate reader. This was done in two independent trials with four independent replicates.

### 3.5 Biofilm Inhibition Assay

A glycerol stock of *Staphylococcus epidermidis* RP62A (ATCC35984) was revived on Congo red agar (CRA) plates and phenotypically characterized to form rough black colonies suggesting its ability to produce the biofilm. A single colony from the CRA plate was transferred to 50 mL trypticasein soy broth supplemented with glucose (TSBg) and grown overnight at 35 °C, 150 rpm. The turbidity of the culture was adjusted to match the 0.5 McFarland standard (1 x 10^8^ cells/mL) and diluted 100-fold prior to inoculation in a 96-well plate. These were treated with compounds **1-7**, in two-fold dilution, starting with 0.125 µg/mL.TSBg served as the negative control, and positive controls were chloramphenicol (8and 4 µg/mL) (Sigma-Aldrich) for planktonic cells and dispersin B (50and 5 µg/mL) (Kane Biotech) for biofilms. The plates were incubated for 20 h at 37 °C without shaking to facilitate biofilm formation. At the end of the incubation period, the planktonic suspension was transferred to another 96-well plate to assess cell viability. An aliquot of 0.02% v/v resazurin (20 µL) was added to each well and incubated for 30 min at 37 °C, 150 rpm. Fluorescence was measured using a 530 nm excitation filter and a 590 nm emission filter. To quantify the formed biofilm, 100 µL-aliquot of wheat germ agglutinin Alexa Fluor 488 (WGA488) probe (50 µg/mL) was added to each well of the original plate and incubated in the dark at 4 °C for 5 h. At the end of incubation, plate was washed with sterile PBS (3x) and air dried at room temperature to remove the unbound WGA488 probe. Bound stain to the biofilm was dissolved in 33% glacial acetic acid and quantified by fluorescence measurement at 485nm/420 nm excitation filter and 528 nm/520 nm emission filters. The minimum inhibitory concentration of compounds causing 100% biofilm inhibition was obtained through CompuSyn software by plotting concentration vs. % cell inhibition. This was done in two independent trials with four independent replicates.

### 3.6 Mammalian Antiproliferative Assay

MDCK NBL-2 (ATCCCCL-34) cells were cultured in Minimum Essential Medium (MEM) supplemented with 10% fetal bovine serum, penicillin (100 units) and streptomycin (100 *µ*g/mL) (Gibco) under a humidified environment with 5% CO_2_ at 37 °C. MDCK (7,500 cells/well) were seeded in 96-well plates and treated after 24 h with varying concentrations of **1-7**, doxorubicin (0.1 and 1*µ*M) and solvent control (1% DMSO). After 72 h, the medium was removed, and 15 *µ*L of 5 mg/mL MTT reagent was added to each well. The plate was incubated for another 3 h at 37 °C with 5% CO_2_ prior to addition of DMSO (100 *µ*L). The absorbance was read at 570 nm using a microplate reader (Biotek Synergy HT). IC_50_values were obtained using Graph Pad Prism 5 based on a 4-point sigmoidal non-linear regression analysis of cell viability vs. log concentration of inhibitor. This was done in two independent trials with four independent replicates.

## Supplementary Materials

Figure S1-S6: HRMS, ^1^H NMR, COSY, HSQC, HMBC and TOCSY spectra of turneroic acid (**1**); Figure S7-S8: LC-ESI-MS of **2** and **3**; Figure S9-S10: MS/MS spectra of **2** and **3**; Figure S11-S12: ^1^H NMR spectra of **2** and **3**.

## Author Contributions

Conceptualization, N.M.L, J.M.R, L.A.S-R and G.P.C.; methodology, N.M.L, C.V.R, J.M.R., M.R.D.P, J.O.T, L.A.S-R; writing – original draft preparation, N.M.L, L.A.S-R and G.P.C.; review and editing - N.M.L, B.W.M., M.G.H., E.W.S., L.A.S-R and G.P.C.; funding acquisition, M.G.H and G.P.C..

