## Supplementary Data for "Inhibition of Biofilm Formation by Modified Oxylipins from the Shipworm Symbiont *Teredinibacter turnerae*"

Contents:

**Table S1.** Minimum inhibitory concentration (MIC) of **1-7** against biofilm formation of *S. epidermidis* RP62A ATCC 35984™ and planktonic cells of *S. epidermidis* RP62A ATCC 35984™, *S. aureus* ATCC 6538™ and methicillin resistant *S. aureus* ATCC 43300™. Selectivity index was computed as a ratio of $MIC_{Planktonic}$/ $MIC_{Biofilm}$.

**Figure S1.** HRMS spectrum of turneroic acid (**1**) $[M+Na]^+$ in positive mode.

**Figure S2.** $^1H$ NMR Spectrum of turneroic acid (**1**) in $CD_3OD$ (500 MHz).

**Figure S3.** COSY Spectrum of turneroic acid (**1**) in $CD_3OD$ (500 MHz).

**Figure S4.** HSQC Spectrum of turneroic acid (**1**) in $CD_3OD$ (500 MHz).

**Figure S5.** HMBC Spectrum of turneroic acid (**1**) in $CD_3OD$ (500 MHz).

**Figure S6.** TOCSY Spectrum of turneroic acid (**1**) in $CD_3OD$ (500 MHz).

**Figure S7.** LC- ESIMS of Compound **2**.

**Figure S8.** LC- ESI-MS of Compound **3.**

**Figure S9.** MS/MS of **2** at 20V in positive mode.

**Figure S10.** MS/MS of **3** at 20V in positive mode.

**Figure S11.** $^1H$ NMR spectrum of **2** in $CD_3OD$ (500 MHz).

**Figure S12.** $^1H$ NMR spectrum of **3** in $CD_3OD$ (500 MHz).

**Table S1.** Minimum inhibitory concentration (MIC) of **1-7** against biofilm formation of *S. epidermidis*RP62A ATCC 35984™ and planktonic cells of *S. epidermidis* RP62A ATCC 35984™, *S. aureus* ATCC 6538™ and methicillin resistant *S. aureus* ATCC 43300™.

| **Compounds** | **MIC (μg/mL) *S. epidermidis* RP62A (ATCC 35984)** | | **Selectivity Index ($MIC_{Planktonic}$/ $MIC_{Biofilm}$)** | **MIC (μg/mL) *S. aureus* (ATCC 6538)** | **MIC (μg/mL) methicillin- resistant *S. aureus* (ATCC 43300)** | **$IC_{50}$ (μg/mL) MDCK NBL-2 (ATCC CCL-34)** |
|---|---|---|---|---|---|---|
| | **Biofilm** | **Planktonic** | | | | |
| **1** | 32.0 | 128 | 4 | >128 | >128 | 61.7 |
| **2** | 90.0 | >128 | >1.4 | >128 | >128 | 33.0 |
| **3** | 11.2 | 24.5 | 2.2 | >128 | >128 | 13.6 |
| **4** | >128 | >128 | ~1 | >128 | >128 | > 128 |
| **5** | >128 | >128 | ~1 | >128 | >128 | > 128 |
| **6** | 16.0 | >128 | >8 | >128 | >128 | > 128 |
| **7** | >128 | > 128 | ~1 | 64.0 | >128 | >128 |
| dispersin B | 50.0 | >>500 | >>10 | *n/a* | *n/a* | *n/a* |
| chloramphenicol | *n/a* | 8 | *n/a* | *n/a* | *n/a* | *n/a* |
| oxacillin | *n/a* | *n/a* | *n/a* | 8 | 1 | *n/a* |

[a] Data presented as MIC of two independent trials performed in quadruplicates. *n/a* = not applicable.

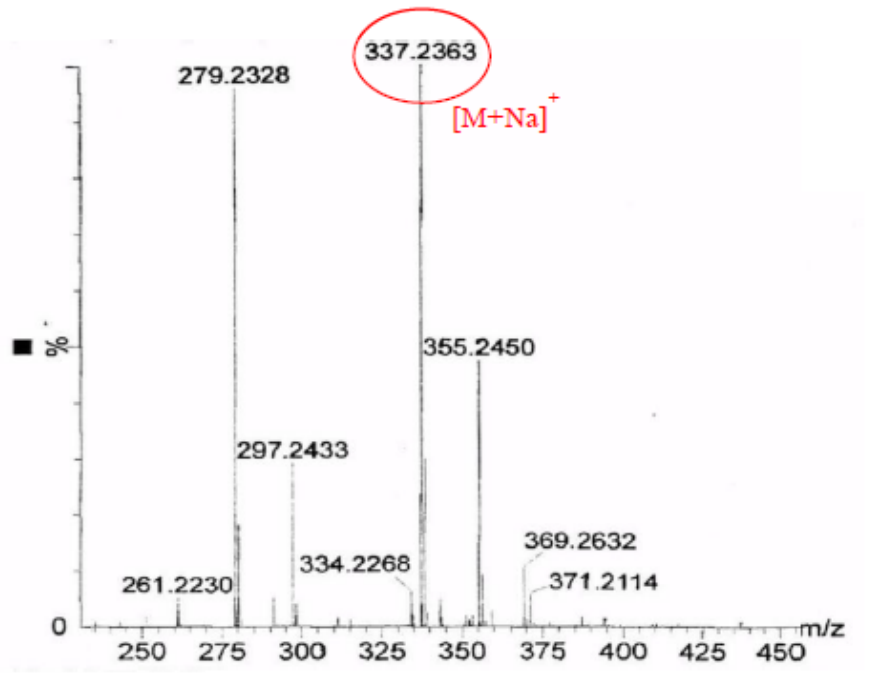


**Figure S1.** HRMS spectrum of turneroic acid (**1**) [M+Na]+ in positive mode.

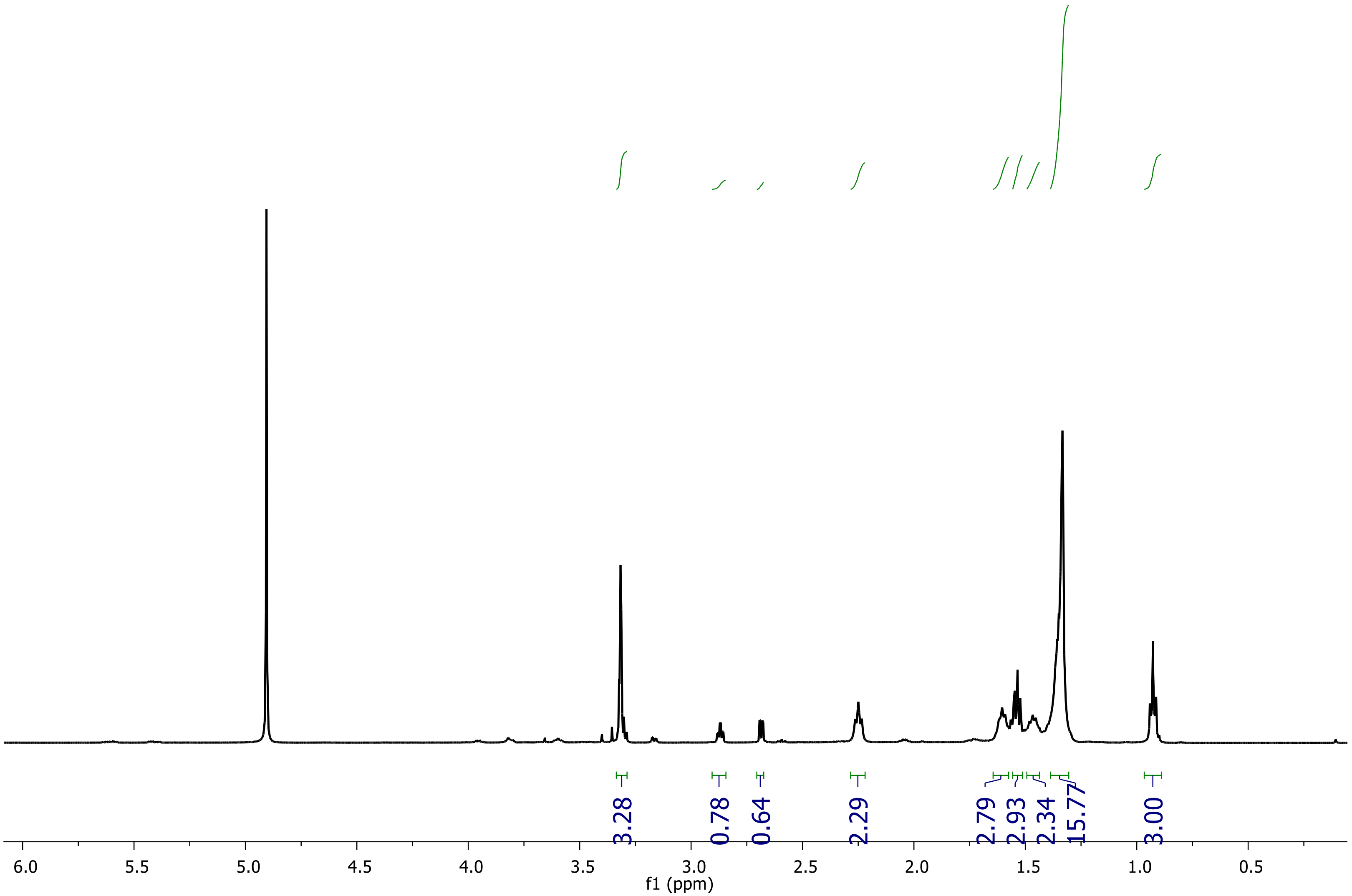


**Figure S2.** $^1$H NMR Spectrum of turneroic acid (**1**) in $CD_3OD$ (500 MHz).

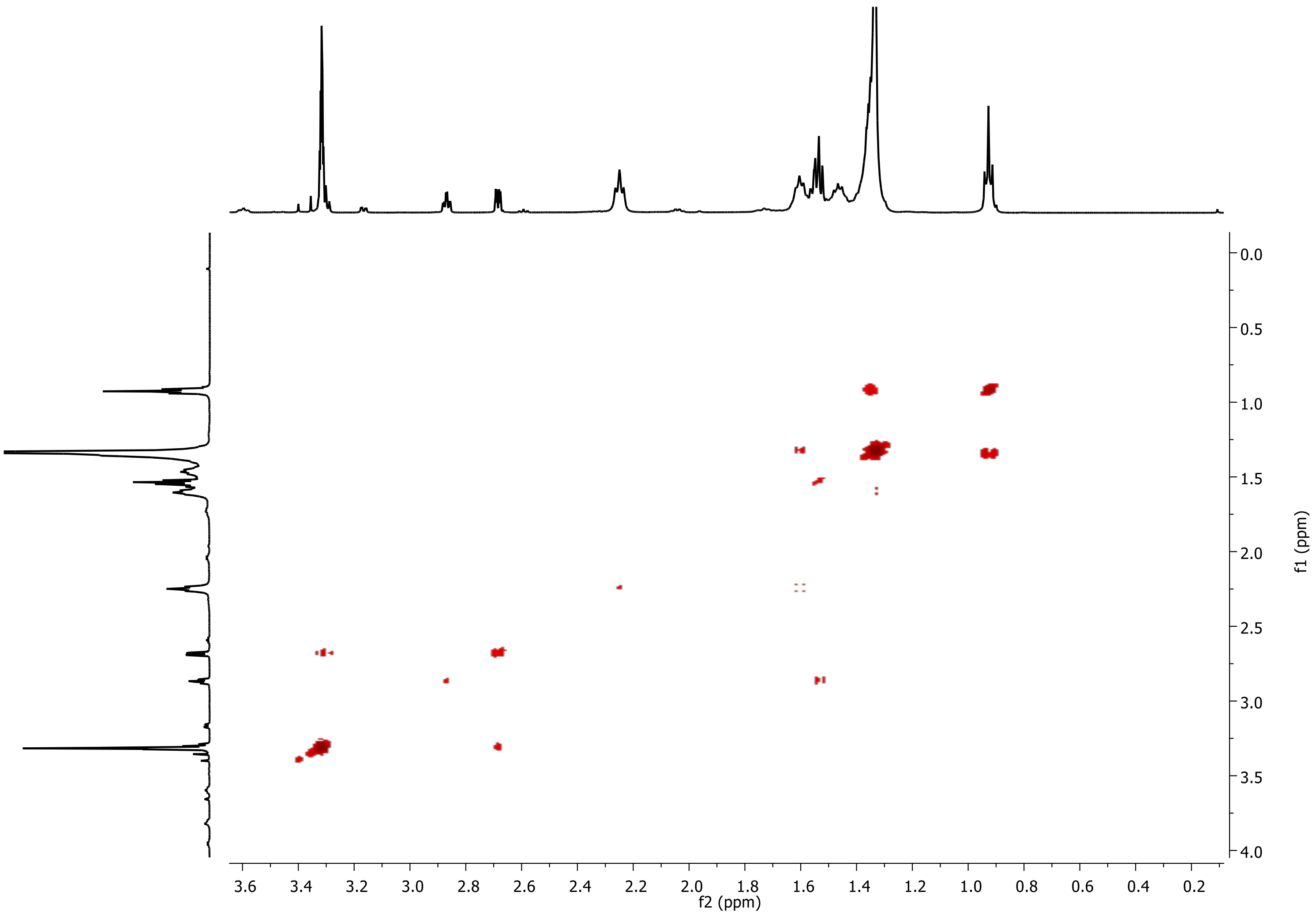


**Figure S3.** COSY Spectrum of turneroic acid (**1**) in $CD_3OD$ (500 MHz).

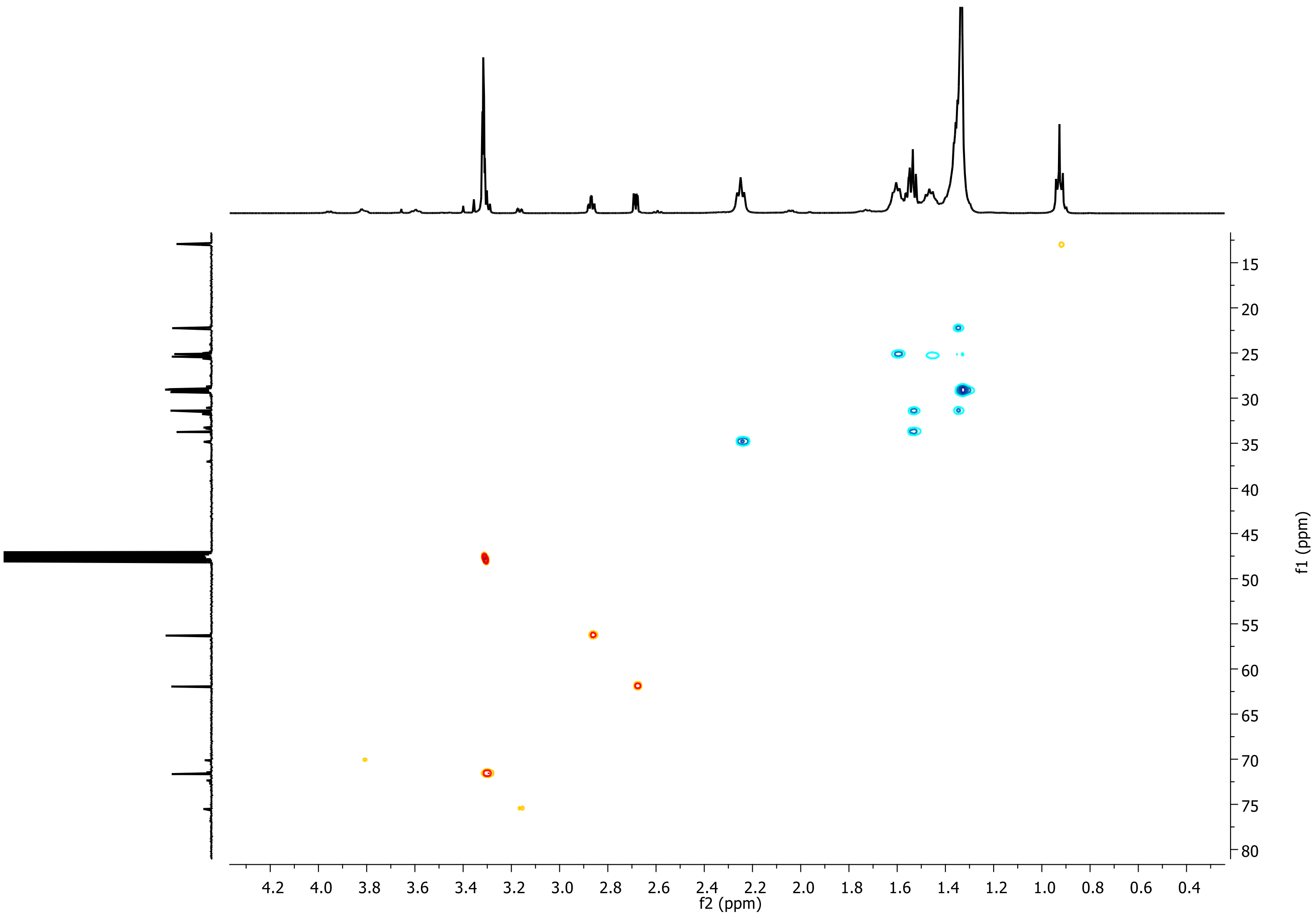


**Figure S4.** HSQC Spectrum of turneroic acid (**1**) in $CD_3OD$ (500 MHz).

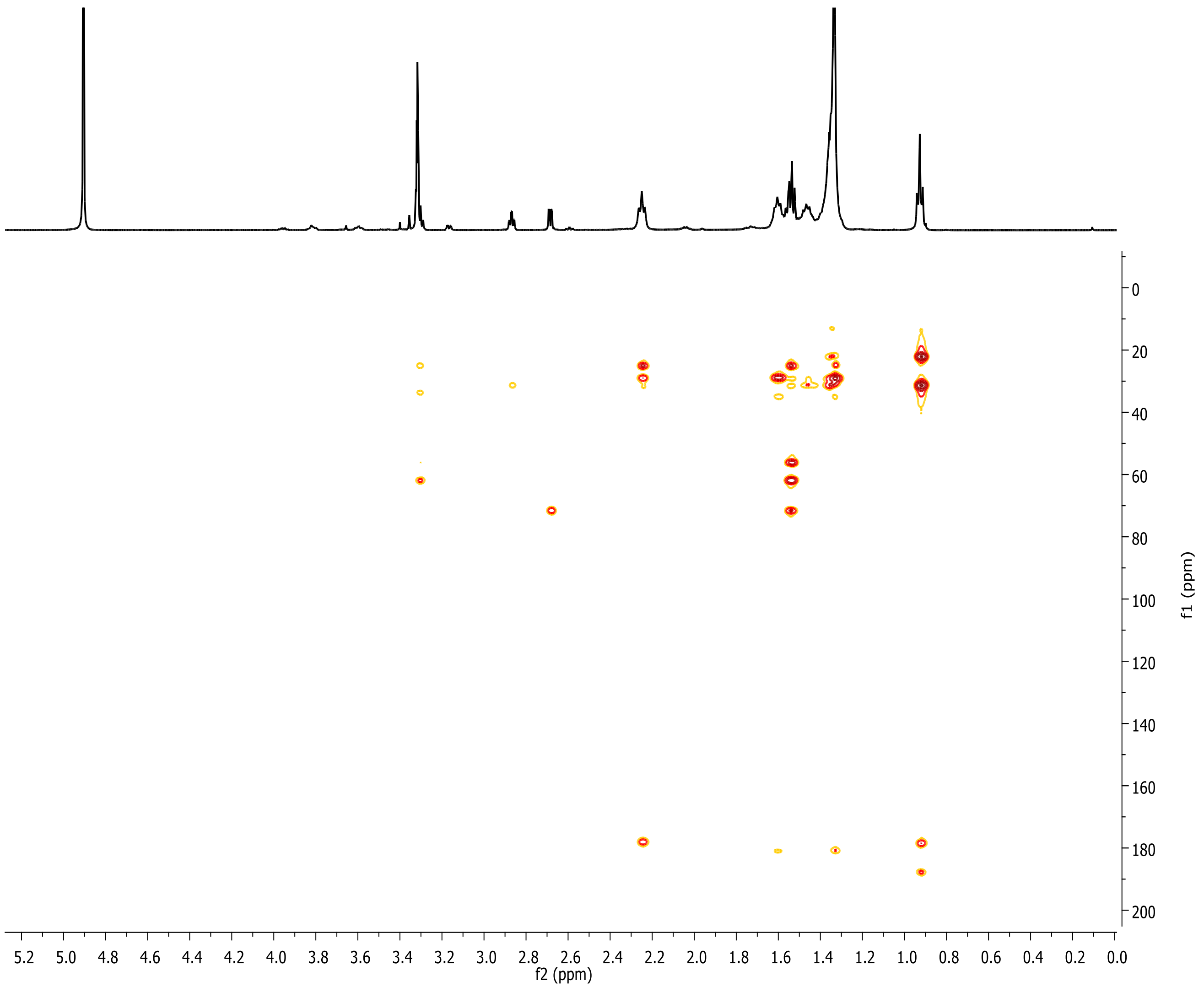


**Figure S5.** HMBC Spectrum of turneroic acid (**1**) in $CD_3OD$ (500 MHz).

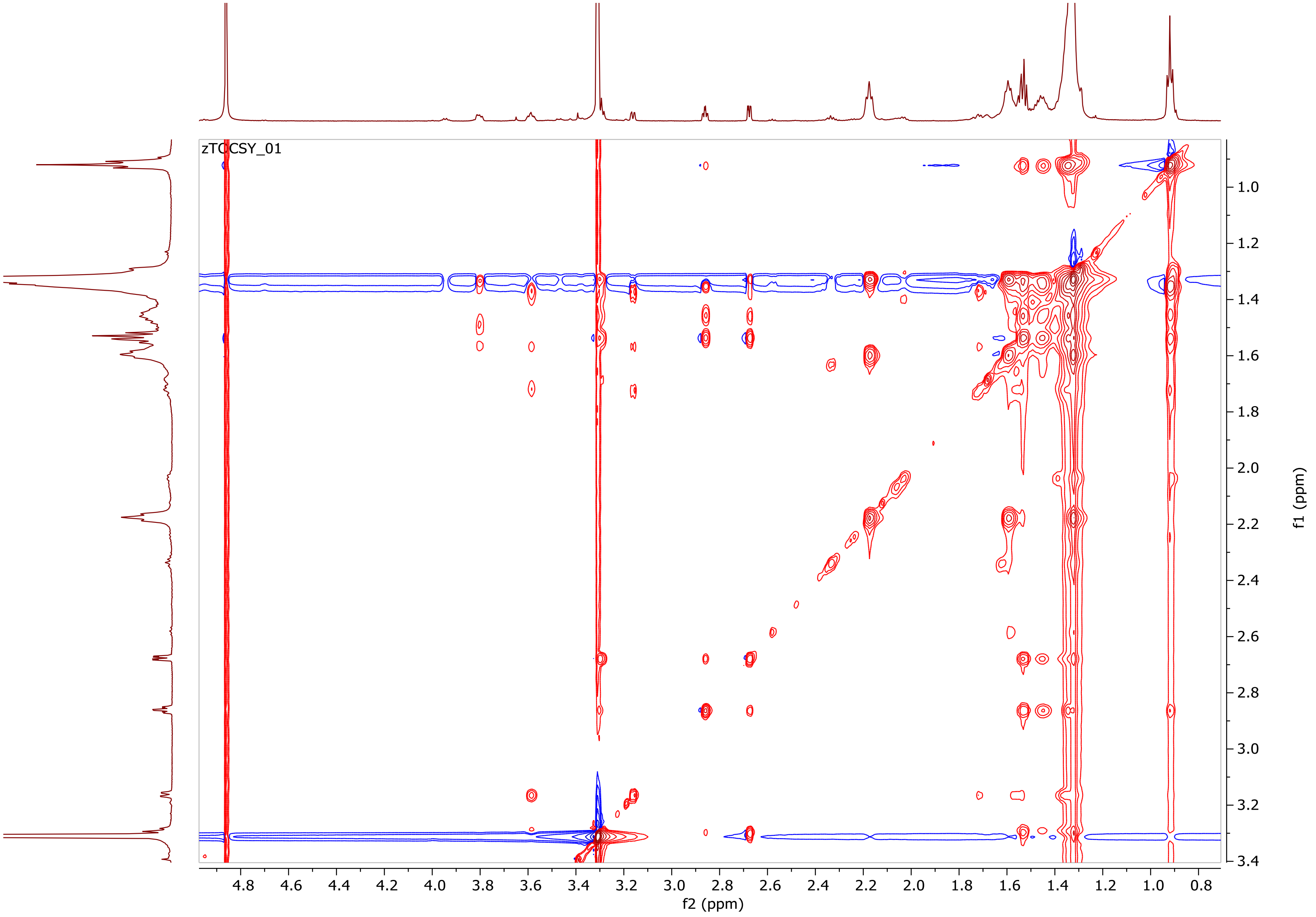


**Figure S6.** TOCSY Spectrum of turneroic acid (**1**) in $CD_3OD$ (500 MHz). The TOCSY spectrum showed the presence of minor components in turneroic acid after prolonged storage in solution.

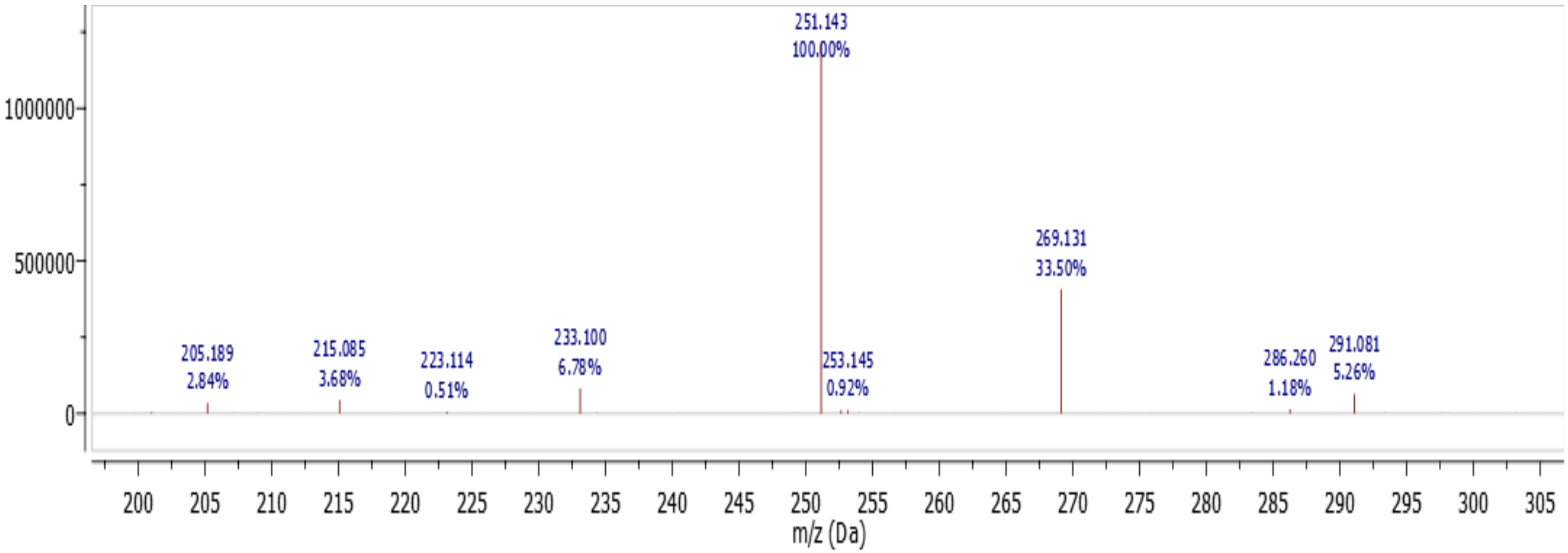


**Figure S7.** LC- ESIMS of **2** $[M+H]^+$ = 269.131, $C_{16}H_{28}O_3$

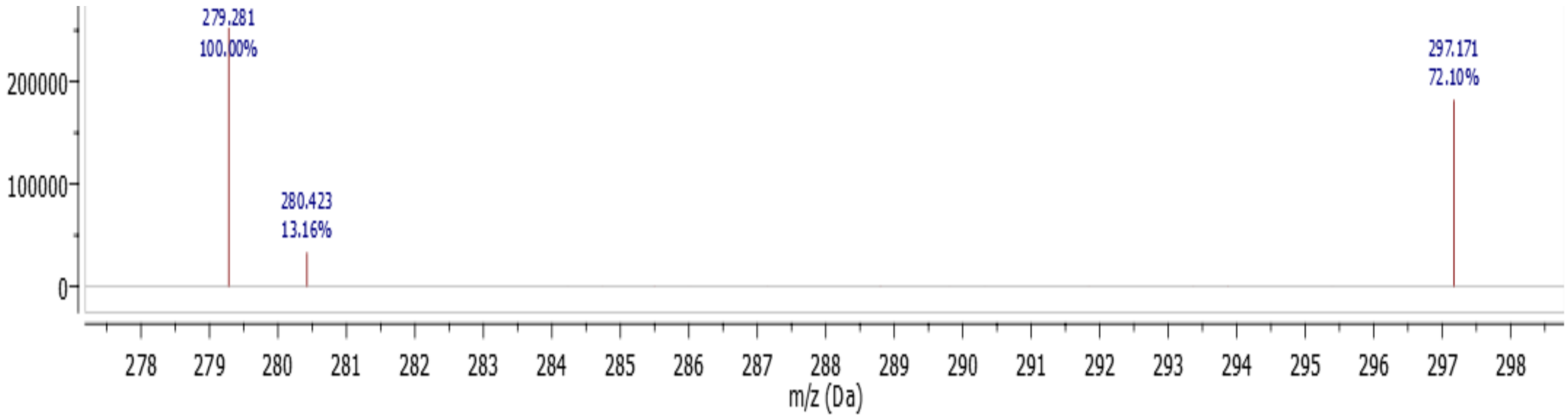


**Figure S8.** LC- ESIMS of **3** $[M+H]^+$ = 297.171, $C_{18}H_{32}O_3$

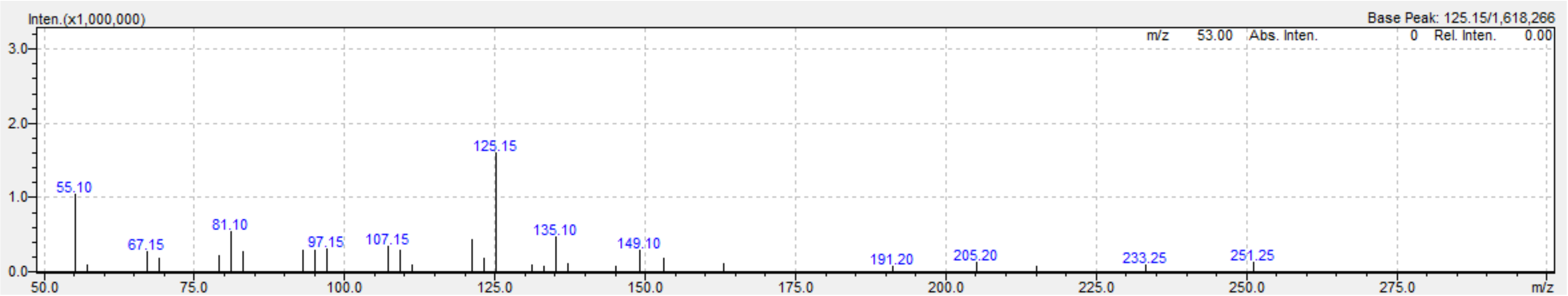


**Figure S9.** MS/MS of **2** at 20V in positive mode.

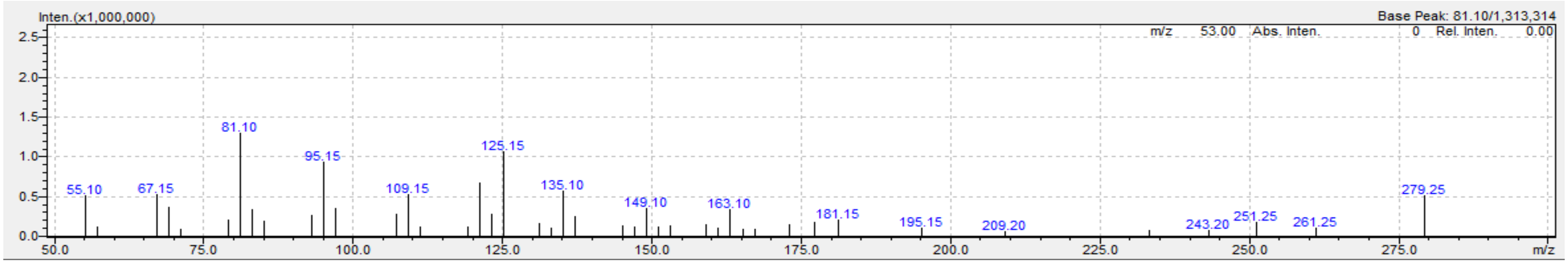


**Figure S10.** MS/MS of **3** at 20V in positive mode.

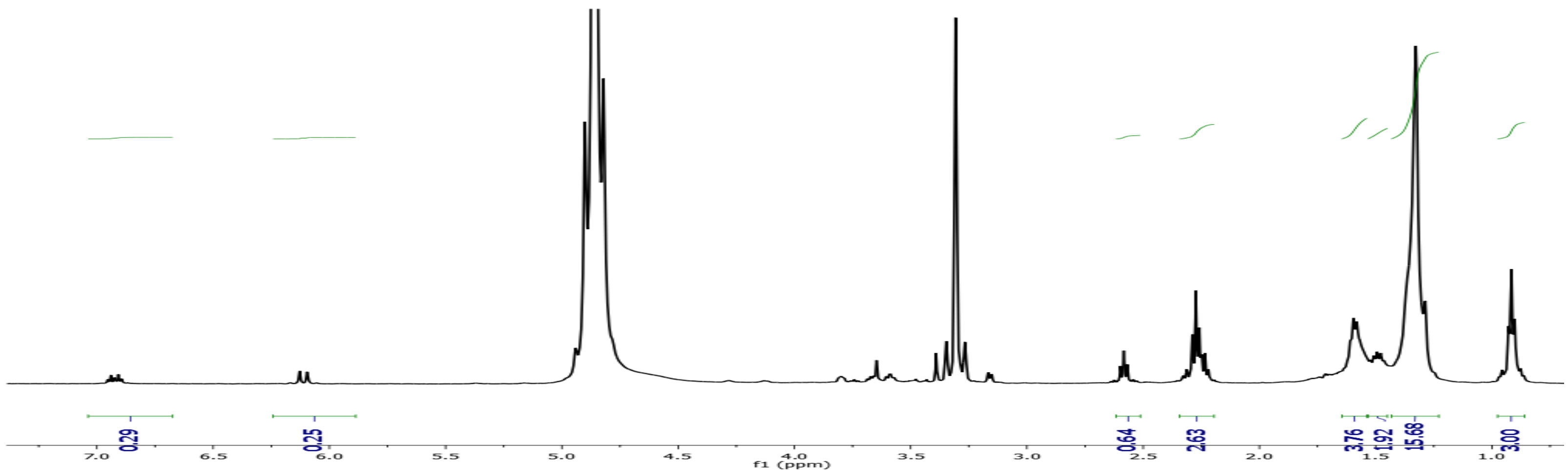


**Figure S11.** [1]H NMR spectrum of **2** in $CD_3OD$ (500 MHz).

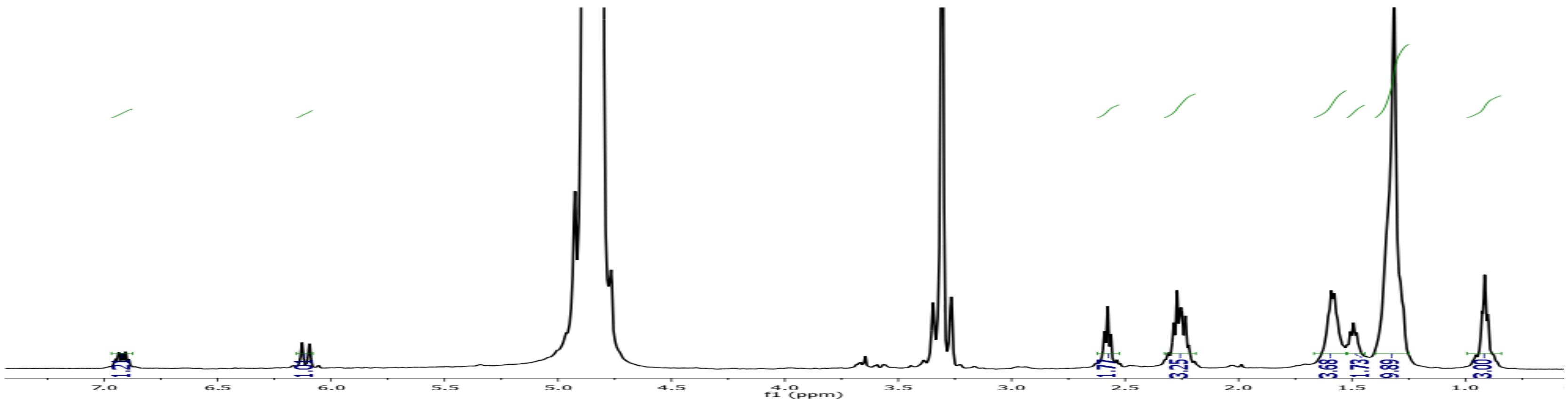


**Figure S12.** [1]H NMR spectrum of **3** in $CD_3OD$ (500 MHz).
